# CRISPR/Cas12a-Mediated Knockout of *INNER NO OUTER (INO)* Gene in *Musa balbisiana* cv. Bhimkol

**DOI:** 10.64898/2026.05.13.724745

**Authors:** Maske Neha Chandrakant, Abhik Gogoi, Dhanawantari L Singha, Seon-Kap Hwang, Thomas W. Okita, Salvinder Singh

## Abstract

Banana (*Musa* spp.) is a vital staple food and cash crop cultivated in over 140 countries, providing nourishment and livelihoods to more than 400 million people worldwide. In this context, Bhimkol (*Musa balbisiana*, BB genome), a diploid banana variety native to Northeast India holds significant nutritional and commercial value. Its high iron and nutrient content have already been commercially validated through products like Bhimvita and Bhimshakti, which utilize fresh fruit pulp as nutrient-rich food for infants. However, Bhimkol fruits typically contain 100–150 seeds, an undesirable trait for product development. The manual removal of these seeds significantly increases production time and labour costs. Furthermore, because bananas are recalcitrant to traditional breeding, there is a constant need for rapid *in vitro* transformation protocols. To address these challenges, as a proof of concept, our research aims to knockout the *INNER NO OUTER* (*INO*) gene, which is responsible for ovule development. Using CRISPR/Cas12a technology, we established an efficient and reproducible *in vitro* regeneration and transformation system using Embryogenic Cell Suspensions (ECS). The resulting CRISPR-edited plantlets exhibited various mutations, including insertions and deletions (INDELs) within the targeted *INO* gene. These INDELs resulted in frameshift mutations that triggered premature stop codons. While these genetic changes are expected to render the banana seedless, phenotypic verification is currently underway to confirm the absence of seeds in mature fruit.

**Significance Statement:** Despite its superior nutritional profile, the commercial viability of the Bhimkol banana (*Musa balbisiana*) is restricted due to abundance of seeds (100–150 per fruit). This study employs CRISPR/Cas12a-mediated knockout the *INNER NO OUTER* (*INO*) gene in Bhimkol and expected to develop seedless fruits. The resulting plantlets exhibit targeted indels that trigger frameshift mutations, effectively disrupting ovule developmental *INO* gene.

## Introduction

Fruit crops are integral to human nutrition and provide essential vitamins, minerals, antioxidants, and fibre. The recommended daily intake of fruits and vegetables is approximately 400 g (Hanke and Flachowsky, 2010). Among these, bananas are the second most cultivated crop globally and the fourth most cultivated crop in developing countries (Moffat, 1999). India, a major centre for Musa diversity, produced approximately 34.5 million tons of bananas in 2022 (FAO, 2022). Diploid banana varieties, including wild species such as *Musa acuminata* and *Musa balbisiana*, exhibit genetic resilience to biotic and abiotic stresses. Hybridization between these species has resulted in various genome compositions, with triploid cultivars such as AAA, AAB, and ABB, which are the most widely consumed (Heslop-Harrison and Schwarzacher, 2007).

Bhimkol, a diploid *M. balbisiana* variety, is predominantly cultivated in Northeast India, particularly in Assam, covering more than 8000 hectares. Nutritionally, Bhimkol contains protein, carbohydrate, dietary fiber, fat and other minerals such as nitrogen, calcium, iron, phosphorus, magnesium and potassium. Bhimkol also reported to possess antibacterial properties, further enhancing its economic significance. The commercial importance of Bhimkol was realized when Bhimkol-fresh pulp derived baby food products, such as Bhimvita and Bhimshakti, have gained market presence beyond Northeast India (Northeast news, April 30, 2014). However, unlike other commercially grown bananas, Bhimkol contains 100-150 seeds per fruit. Possession of many seeds is considered as undesirable trait as more seeds mean less pulp. Further consumption of seeds in the form of processed powder may cause digestive issues in infants. Removal of seeds prior to large-scale processing will add an extra labour-intensive step, thereby increasing the overall cost of production. Further, with changing market preferences and urban lifestyles, seedless fruits are emerging as the preferred choice (Premachandran *et al*., 2019). Therefore, development of seedless Bhimkol variety is need of the hour. Techniques such as stenocarpy and parthenocarpy are not very effective in *Musa balbisiana* cv. Bhimkol because the plant has strong genetic mechanism controlling seed development. As a result, fertilization does not fully occur (incomplete fertilization), which limits the success of these techniques. To overcome these challenges, application of advanced, precise approaches such as gene-editing technologies offers a promising alternative. For seedless fruit development, genes responsible for ovule formation, such as the *INNER NO OUTER* (*INO*) gene, a member of the YABBY family that is crucial for outer integument development, can be potential targets for precise gene-editing techniques (Baker *et al*., 1997; Villanueva *et al*., 1999). Mutations in the *INO* gene, as observed in *Arabidopsis* and *Annona squamosa*, lead to incomplete ovule formation, and consequently, seedlessness (Baker *et al*., 1997; Gaiser *et al*., 1995). A naturally occurring mutation in *A. squamosa* resulted in a seedless Thai variety, which was attributed to deletion of the *INO* gene (Lora *et al*., 2011). Rodrigues *et al*., (2024) demonstrated that the presence or absence of seeds in *A. squamosa* is controlled by a single fully recessive gene, with seedless phenotypes arising from deletion of the *INO* locus. Given that seedlessness is linked to genetic mutations that affect ovule integrity, genome editing of the *INO* gene in Bhimkol could potentially yield seedless varieties with desirable traits.

Advanced genome editing technique such as CRISPR/Cas12a system has recently emerged as a superior alternative to CRISPR/Cas9, offering an advanced and efficient genome-editing tool (Bin Moon *et al*., 2018). The Cas12a endonuclease belongs to the Class 2, Type V CRISPR system and is derived from bacterial species such as *Prevotella* and *Francisella*. Among the three Cas12a homologs, FnCas12a (*Francisella novicida*), LbCas12a (*Lachnospiraceae bacterium*), and AsCas12a (*Acidaminococcus sp*.) have been extensively utilized in plant genome-editing applications (Zetsche *et al*., 2015). Cas12a is characterized by a smaller protein size than Cas9 and requires a shorter CRISPR RNA (crRNA) for effective function (Liu *et al*., 2017). Unlike Cas9, which relies on a dual-RNA complex, Cas12a is guided by a single crRNA molecule. It specifically recognizes a thymine-rich protospacer adjacent motif (PAM) located upstream of the target site and induces DNA cleavage through staggered cuts, generating 5-base pair (bp) overhangs (Zetsche *et al*., 2015). Structurally, Cas12a possesses only a single endonuclease domain, RuvC (NUC lobe), which catalyses the sequential cleavage of the non-target DNA strand, followed by the target strand (Yamano *et al*., 2016). The resulting staggered ends are advantageous for non-homologous end-joining (NHEJ)-mediated gene insertion, particularly in mammalian genomes (Maresca *et al*., 2013). This feature is especially beneficial for genome editing in non-dividing cells, where homology-directed repair (HDR) is inefficient (Chan *et al*., 2011). The CRISPR/Cas12a system enables gene deletion, insertion, base editing, and locus tagging in monocots and dicots, with minimal off-target effects. Its versatility and ease of use have increased its widespread adoption. The effectiveness of this system has been demonstrated in a range of crop species including tobacco, rice, soybean, and wheat (Kaur *et al*., 2020; Bandyopadhyay *et al*., 2020).

Khanna *et al*., (2004) developed a highly efficient method for the *Agrobacterium*-mediated transformation of banana embryogenic cell suspensions (ECS). This system enabled the generation of transformants with notable efficiency from cultivars belonging to the commercially significant AAB and AAA genomic groups. ECS has also been established in cultivars using meristems (Xu *et al*., 2005; Kulkarni *et al*., 2006; Strosse *et al*., 2006) or immature flowers (Côte *et al*., 1996; Grapin *et al*., 1996; Ganapathi *et al*., 2001; Khalil *et al*., 2002). However, male flowers showed the highest responsiveness to initiating embryogenic culture in the Grand Nain cultivar (Escalant *et al*., 1994; Côte *et al*., 1996; Navarro *et al*., 1997; Sági *et al*., 1998; Becker *et al*., 2000). The present study aimed to utilize CRISPR/Cas12a as a proof of concept to knock out the *INO* gene in Bhimkol banana cultivar. It is expected that these mutants will exhibit defective ovules, reduced seed numbers, or completely seedless fruit. Developing a seedless Bhimkol variety would significantly enhance its commercial viability, effectively meeting both consumer and industrial demands.

## Material and Methods

### Guide RNA design and CRISPR/Cas12a construction

The *INO* gene from Bhimkol (1175 bp), which was previously isolated, cloned, and sequenced (Gogoi, 2018), was used to design sgRNA1. The 23-nucleotide sgRNA1 was designed to target exon 2 of *INO* gene using Benchling (https://benchling.com), with selection based on off-target scores. This sgRNA1 sequence was cloned into the expression vector pSH924A, which is regulated by the rice *OsU6B* promoter and Pol III terminator. For cloning into pSH924A, 0.5 µmoles each of forward (sgRNA1-F, 5’AGATGGTGAGTGTTCCATGCAGCAGCC3’) and reverse (sgRNA1-R, 5’AAAAGGCTGCTGCATGGAACACTCACC3’) oligonucleotide primers incorporating AGAT and AAAA adapter sequences, respectively, were boiled for 20 min, annealed, and ligated with 1 µg of the *Bsa*I-digested pSH924A. Following ligation, transformed colonies were identified by polymerase chain reaction (PCR) amplification of the expression cassette (U6B2::INO-sgRNA1::Pol III) using U6B2-*Bsa*I forward (5’AAGGTCTCTGGCACGATCTGCCGCCGGATCATG3’) and U6B2t-*Bsa*I reverse (5’AGAGGTCTCAAAACTGACCATTTGTGGGTCTGT3’) primers, producing a 457 bp fragment with *Bsa*I recognition sites at both ends of the fragment. PCR was performed with an initial denaturation at 95 °C for 4 min, followed by 35 cycles of 95 °C for 30 s, 56 °C for 30 s, and 72 °C for 1 min, and a final extension at 72 °C for 5 min. The PCR product containing the expression cassette was subsequently digested with *Bsa*I and cloned into the pSH916 binary vector. Colony PCR using M13RV (5’TTTCACACAGGAAACAGCTATGAC3’) and 916seqR2 (5’AATCGCGCACGTACGGTTGAC3’) primers was performed and the PCR products were analysed on a 0.8% agarose gel. DNA sequencing was performed to confirm the successful cloning of the sgRNA1 expression cassette into the pSH916 vector. The validated construct, designated as pSS03, was used for the genetic transformation of Bhimkol mediated through *Agrobacterium tumefaciens* using strain AGL1.

### Delivery of the assembled binary construct to Bhimkol

An embryogenic cell suspension (ECS), derived from the embryogenic calli of *Musa balbisiana* (Bhimkol, BB genome), was utilized for *Agrobacterium*-mediated genetic transformation. Immature male flowers (comprising up to 15 bracts) served as explants to induce embryogenic calli, which facilitated the establishment of the ECS (Longkumer, 2022). The *Agrobacterium* inoculum was prepared in MGL broth supplemented with 50 mg/L kanamycin and 10 mg/L rifampicin, then incubated at 28°C with shaking until an optical density (OD_600_) of 0.6–0.8 was reached. *Agrobacterium* cultures were then harvested and resuspended in *Agrobacterium* resuspension (AA) medium containing 200 μM acetosyringone, followed by a 15-minute incubation with intermittent shaking. The *Agrobacterium* suspension was mixed with the ECS (5 mL packed cell volume; PCV) and incubated for 15 minutes at room temperature. The ECS–*Agrobacterium* mixture was then vacuum-dried on sterile filter paper, transferred to a co-cultivation medium supplemented with 200 μM acetosyringone, and incubated in the dark at 26°C for three days. Following co-cultivation, the infected ECS was washed sequentially with sterile distilled water, with a final rinse containing 250 mg/L timentin. The treated cells were cultured in M3 somatic embryo maturation medium supplemented with 40 mg/L hygromycin and 250 mg/L timentin. These were maintained in the dark at 26°C and sub-cultured at 3 weeks intervals for three rounds of selection. Matured somatic embryos were subsequently transferred to M4 somatic embryo germination medium containing 40 mg/L hygromycin and 250 mg/L timentin. The cultures were maintained under a 16-hour light (7,000 lux) and 8-hour dark photoperiod at 26°C with 80% relative humidity. Finally, well-grown, putative transformed shoots regenerated *in vitro* were transferred to R1 rooting medium (40 mg/L hygromycin and 250 mg/L timentin) and maintained under the same conditions. To eliminate the possibility of “escapes” or false positives, hygromycin and timentin were maintained throughout every stage of the regeneration process. Detailed protocols and media compositions are provided in Supplementary Tables 1–13.

### Molecular analysis of the putative CRISPR-edited Bhimkol plantlets

Genomic DNA was isolated from leaf tissues of both putative CRISPR-edited and wild-type (WT) plants using the cetyltrimethylammonium bromide (CTAB) method (Doyle and Doyle, 1987). Genomic DNA was subjected to PCR using gene-specific primers sgF (5’TCCACTCAATCGCCTGAACT3’) and sgR (5’TGTCTCGTTCTCTGCCTCTC3’) to amplify a 591 bp fragment containing the sgRNA1 target sequence. The PCR conditions included initial denaturation at 95°C for 5 min, followed by 35 cycles of denaturation at 95°C for 30 s, annealing at 55°C for 30 s, and extension at 72°C for 1 min, with a final extension at 72°C for 10 min. The PCR products were analysed using 1.0% agarose gel electrophoresis and documented.

### Nucleotide sequencing and analysis of the sequence results for CRISPR-edited Bhimkol plantlets

The 591 bp amplicons obtained from CRISPR-edited Bhimkol plants were sequenced externally by Eurofins Genomics India Pvt. Ltd., Bengaluru, utilizing Seq-SgF (5’CCTCCCTCCCTCCCCTATAT3’) and Seq-SgR (5’CTGCATGCTTTGGGTCCC3’) primers. Nucleotide sequence analysis was conducted using FASTA sequence alignment and the ICE Synthego tool (https://www.synthego.com/products/bioinformatics/analysis) to detect insertions or deletions (INDELs) within the targeted sgRNA1 region of the *INO* gene in Bhimkol.

### Determination of Amino acid residues and Open Reading Frame of the *INO* gene in the CRISPR-edited Bhimkol plantlets

After confirming successful genomic integration, we evaluated the potential impact at the protein level by examining the amino acid residues and open reading frames (ORFs). The coding sequences (CDS) of the *INO* gene from both wild-type (WT) and CRISPR-edited plants were analyzed using six-frame translation (https://www.bioline.com/media/calculator/01_13.html) and ORF finder tools (https://www.ncbi.nlm.nih.gov/orffinder/) to identify ORFs and translate nucleotide sequences. Protein sequence analysis was performed using SMART (http://smart.embl-heidelberg.de/)to investigate the protein domain architecture between the WT and CRISPR-edited Bhimkol plants. The protein domain structure illustrated by DOG.1 (Ren *et al*., 2009).

## Results

### Design and Validation of sgRNA and CRISPR/Cas12a Constructs

The identified sgRNA1 was optimal based on its low off-target potential and the presence of a T-rich protospacer adjacent motif (PAM), TTTA. For the initial cloning, the sgRNA1 was placed between the *OsU6B* promoter and a *Pol III* terminator of the pSH924A plasmid, which carries an ampicillin resistance gene (Supplementary Figure 1). Successful clones were identified by the absence of *Nco*I digestion, confirming that the sgRNA1 insertion had replaced the *Nco*I site previously flanked by *Bsa*I sites. The resulting *U6B2::sgRNA1::Pol III T* expression cassette (457 bp) was then amplified, digested with *Bsa*I, purified, and ligated into the pSH916 binary vector (Fig. 1A). The transformed ligation mixture into *E. coli* XL10 gold strain yielding a 305 bp amplicon after colony PCR (Fig. 1B). Positive clones were further validated through nucleotide sequencing (Fig. 1C and Supplementary Figure 2). The final confirmed recombinant plasmid was designated as pSS03.

**Figure 1.**
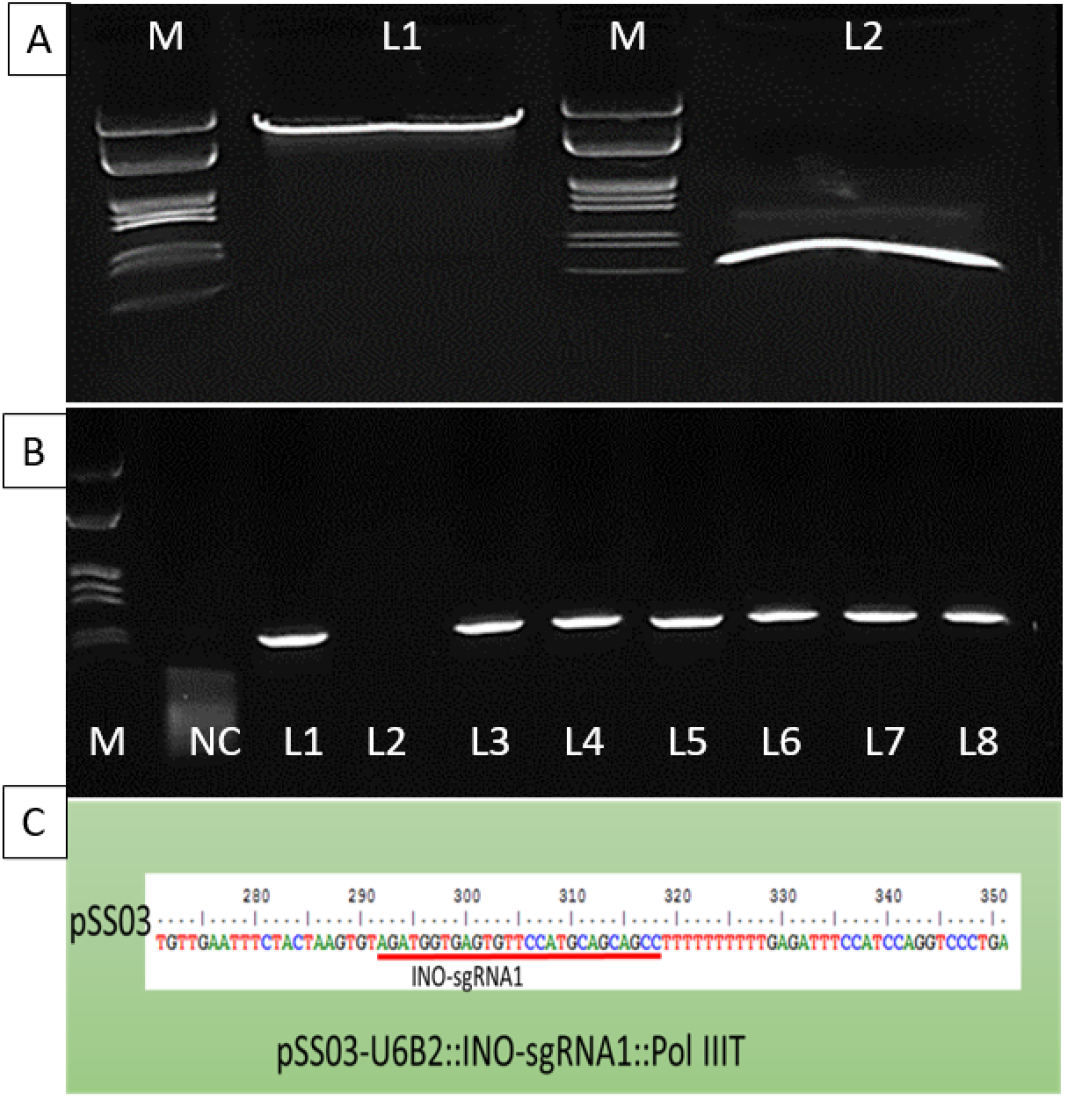
A: Cloning of whole expression cassette U6B2::INO-sgRNA1::Pol III T into pSH916 binary vector. M: 1-Kb Marker; Lane 1: pSH916 digested with *Bsa*I; Lane 2: PCR amplified whole expression cassette, U6B2::sgRNA1::Pol III T digested with *Bsa*I. B: Colony PCR of eight randomly selected colonies to confirm the cloning of whole expression cassette (U6B2::INO-sgRNA1::Pol III T) into pSH916 binary vector. NC: Negative control; Lanes: 1-8: Colony PCR using M13RV forward and 916seqR2 reverse primers. C: Confirmation of the whole expression cassette (U6B2::sgRNA1::Pol III T) into pSH916 (pSS03) through nucleotide sequencing.

### Development of putative CRISPR/Cas12a edited Bhimkol lines

The protocol employed for the development of the embryogenic calli from male floral tissue of Bhimkol was efficient showing numerous numbers of viable embryogenic calli (Fig 2A&B). Subsequently, the embryogenic cell suspension (ECS) developed from the embryonic calli of *Musa balbisiana* cv. Bhimkol was infected with the *Agrobacterium* strain AGL1 containing the CRISPR construct pSS03 (Fig 2C-E). Following the selection rounds, putatively transformed somatic embryos survived and proliferated, while untransformed cells underwent necrosis. Within four weeks, the mature somatic embryos developed green embryoids, which differentiated into shoots within two months. Healthy shoots regenerated in the M4 medium containing 40 mg/L hygromycin, and 250 mg/L timentin after one month of culture (Fig. F). A well-developed rooting was observed from *in vitro* regenerated shoots in the R1 rooting medium containing 40 mg/L hygromycin, and 250 mg/L timentin after 14 days of culture (Fig. G). Putative CRISPR-edited Bhimkol plantlets with healthy shoots and roots, ready for hardening (Fig. H).

**Figure 2.**
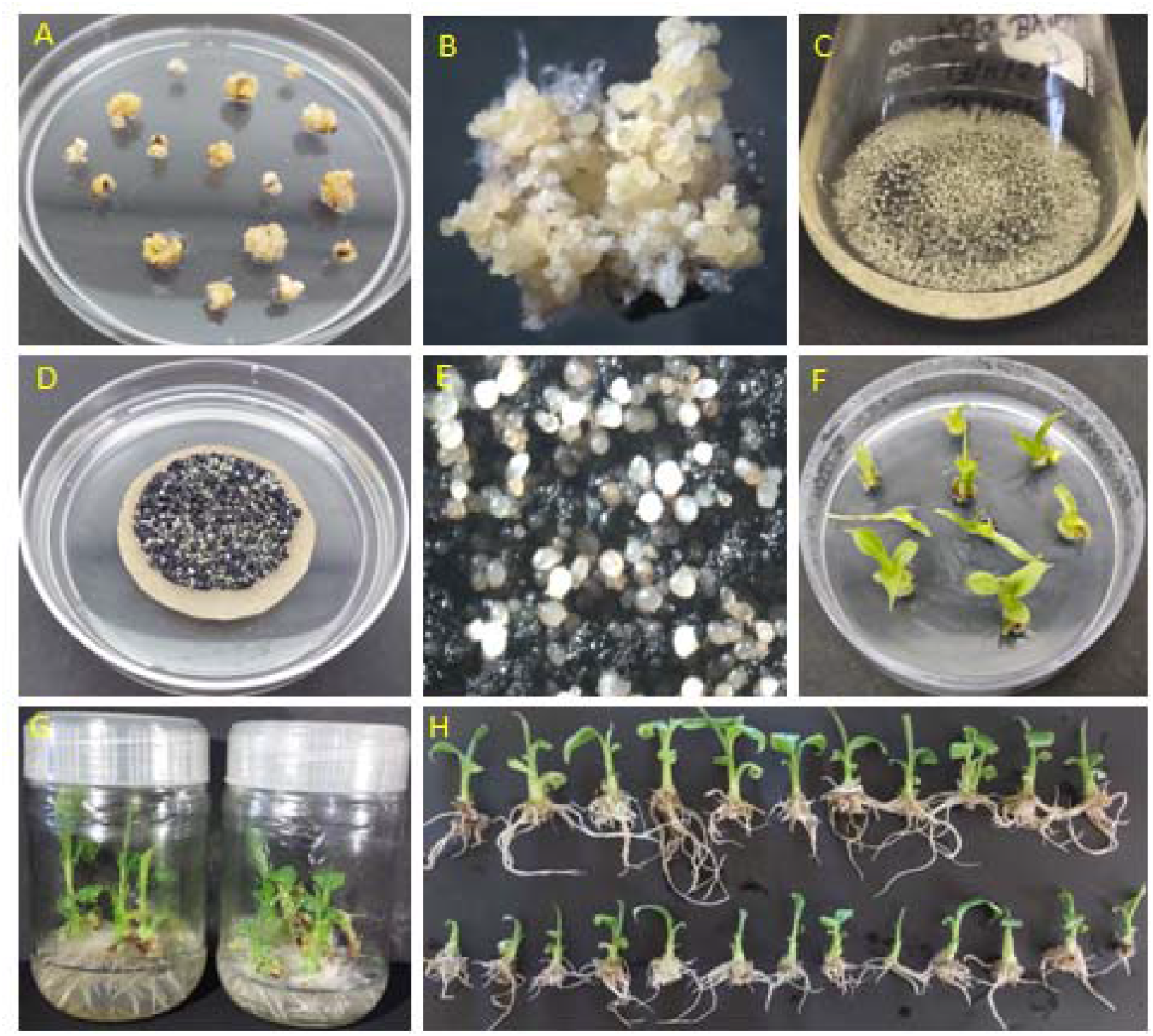
Genetic transformation of Bhimkol mediated through *Agrobacterium* using ECS as explant. A) Induction of embryonic calli from male flowers of Bhimkol after 6 months of culture. B) A magnified view at 40x of embryogenic calli. C) Establishment of ECS after 8 months of culture was used for genetic transformation mediated through *Agrobacterium* harboring pSS03. D) Embryonic calli in selection medium with 40 mg/L hygromycin, and 250 mg/L timentin. E) A magnified view of putative transformed mature somatic embryos at 40x magnification. F) Shoots regeneration in M4 medium containing 40 mg/L hygromycin, and 250 mg/L timentin after one month of culture. G) Rooting from i*n vitro* regenerated shoots in the R1 rooting medium containing 40 mg/L hygromycin, and 250 mg/L timentin after 14 days of culture. H) Putative CRISPR-edited Bhimkol plantlets with healthy shoots and roots, ready for hardening.

### Molecular analysis of CRISPR/Cas12a-edited mutants of Bhimkol

The analysis of the nucleotide sequence result allowed for the identification of insertions and deletions (INDELs) within the *sgRNA1* target site of the *INO* gene in Bhimkol. Multiple sequence alignment revealed that one putative CRISPR-edited plantlet, designated 03_01, exhibited a single base pair (G) insertion at position 144 bp, downstream of the *sgRNA1* site; this mutation was categorized as Type I (Fig. 3A). Another line, 03_05, displayed a 7-bp deletion within the *sgRNA1* target region, spanning positions 97–103 bp (Fig. 3B). Furthermore, analysis using ICE (Inference of CRISPR Edits) software from Synthego showed that lines 03_06 and 03_17 harboured 4-bp deletions within the target region at positions 97–100 and 96–99, respectively. These were classified as Type II mutations (Fig. 3C–D). The overall mutation frequency (Types I and II) in the targeted *INO* gene was calculated to be 23.5%. Despite these genetic modifications, the growth patterns and morphology of the edited lines (03_01, 03_05, 03_06, and 03_17) appeared phenotypically normal compared to the wild type (Fig. 4).

**Figure 3:**
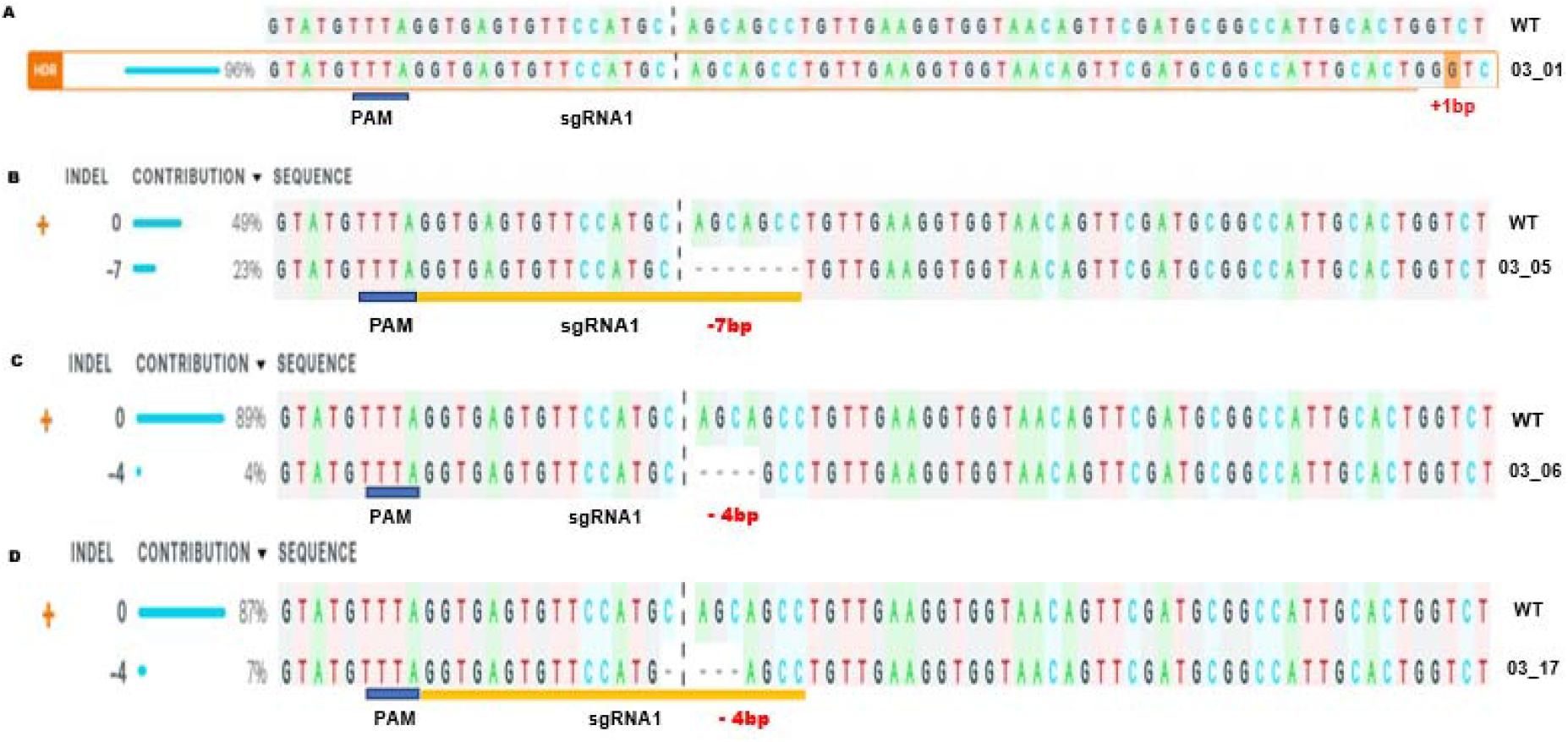
Sequencing result of the CRISPR/Cas12a-edited mutants of Bhimkol. Type-I Mutation A) Insertion of 1 nucleotide (G) downstream of PAM and sgRNA1 sequence at position 144bp in the *INO* gene of Bhimkol. Type-II Mutation B) 03_05 showed deletion of 7 bp downstream of PAM sequence and within sgRNA1 at position ranging from 97 bp to 103 bp, C) 03_06 showed deletion -4bp downstream of PAM sequence and within sgRNA1 at position ranging from 97 bp to 100 bp and D) 03_17 showed deletion of 4 bp downstream of PAM sequence and within sgRNA1at position ranging from 96 bp to 99 bp in *INO* gene of Bhimkol.

**Figure 4:**
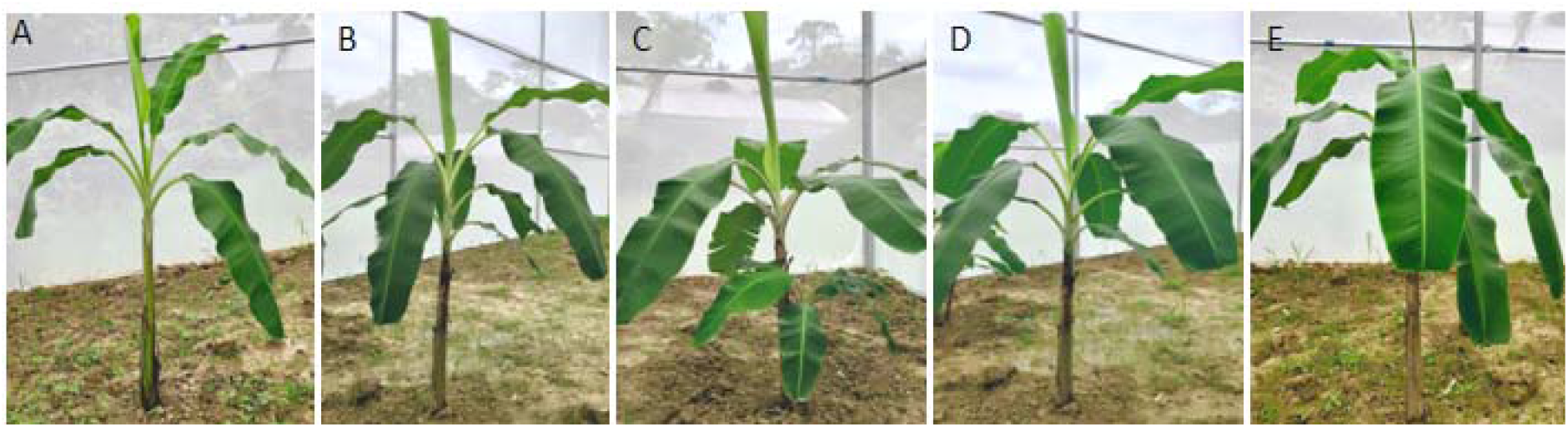
CRISPR-edited Bhimkol plants transferred to the transgenic net house, A) 03_01, B) 03_05, C) 03_06, D) 03_17 and E) Wild Type (Control)

### Determination of Amino acid residues and ORF of the *INO* gene in the CRISPR/Cas12a-edited Bhimkol plantlets

The coding sequences (CDS) of the *INO* gene in wild-type (WT) and CRISPR-edited plants were analyzed using six-frame translation and open reading frame (ORF) finder tools. The WT *INO* gene consists of 645 bp and encodes a 214-amino acid protein in the +1 ORF. In the CRISPR-edited line 03_01, a single-nucleotide (G) insertion caused a frameshift mutation, leading to a premature stop codon at 157 bp and resulting in a truncated 52-amino acid protein. Similarly, lines 03_05, 03_06, and 03_17 also developed premature stop codons at positions 100 bp, 103 bp, and 94 bp, respectively. These mutations produced truncated proteins consisting of 33, 34, and 31 amino acid residues, respectively. Overall, the CRISPR-induced INDELs significantly altered the ORF compared to the WT, leading to premature termination and the production of truncated proteins (Fig. 5).

**Figure 5:**
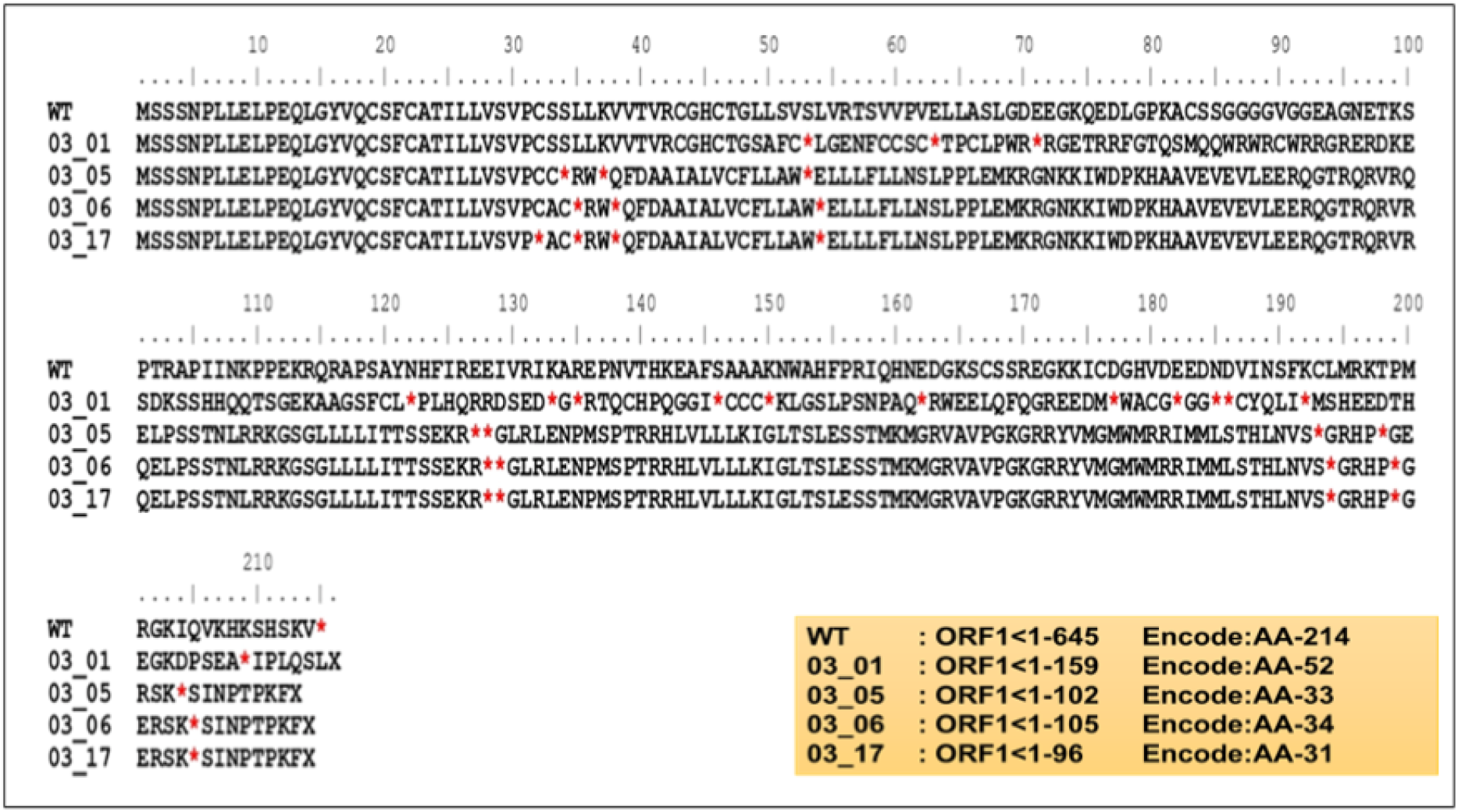
Sequence analysis of the *INO* gene in CRISPR/Cas12a-edited lines. Amino acid sequences from lines 03_01, 03_05, 03_06, and 03_17 were aligned with the wild type sequence. Red asterisks indicate stop codons.

### The protein domains of CRISPR/Cas12a-edited plants were annotated as follows

Protein sequences from wild-type (WT) and CRISPR-edited plants were analysed using the SMART platform for domain annotation and architectural analysis. In WT plants, two distinct protein domains were identified: a YABBY domain and an HMG_BOX 2 domain. The YABBY domain spans residues 9-162 while the HMG_BOX 2 domain is located between residues 111 and 165. In contrast, the CRISPR-edited lines exhibited a complete loss of the HMG_BOX 2 domain. Furthermore, the YABBY domain was severely truncated in all edited lines: 03_01 (residues 9-51), 03_05 (residues 9-33), 03_06 (residues 9-34), and 03_17 (residues 9-31). These alterations demonstrate that CRISPR-mediated INDELs significantly reduced the length of the YABBY domain and eliminated the entire HMG_BOX 2 domain compared to the WT protein architecture (Fig. 6).

**Figure 6:**
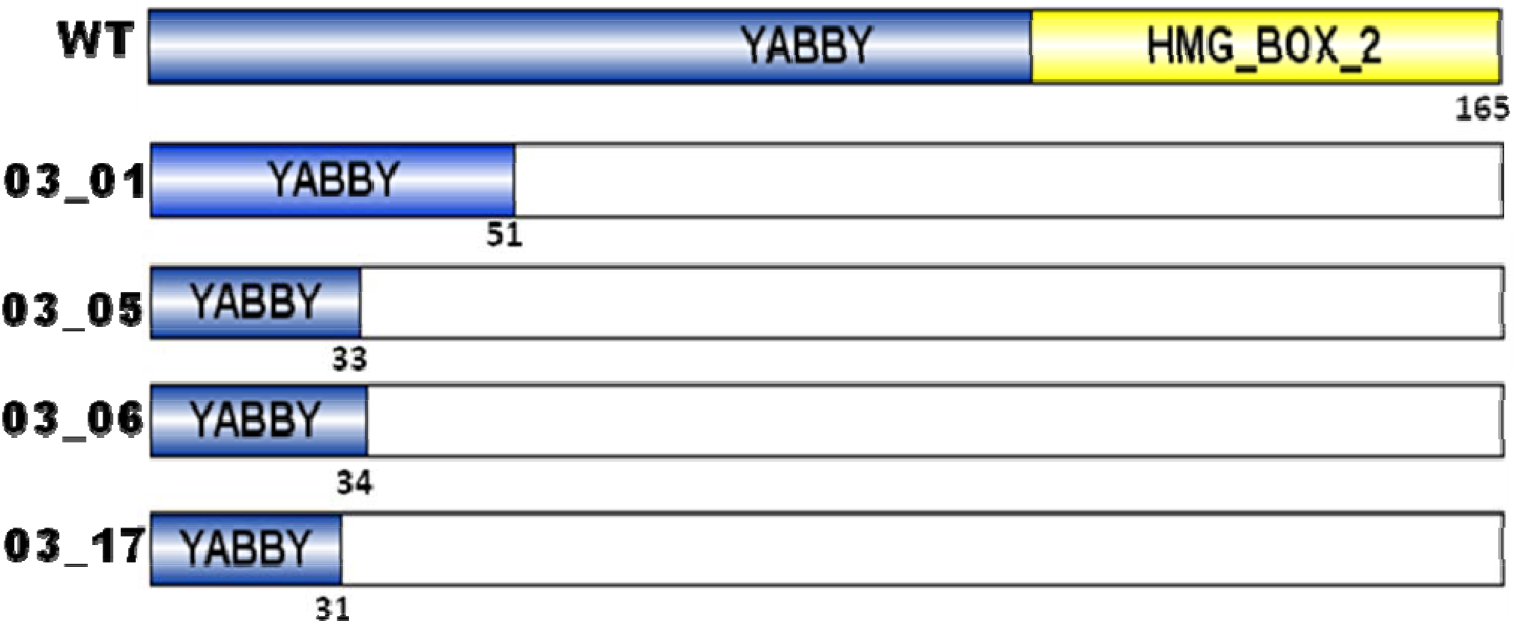
Analysis of YABBY and HMG-box_2 domain in the WT and CRISPR-edited lines. The CRISPR-edited Bhimkol lines exhibit a complete absence of the HMG_box_2 and truncated YABBY domains.

## Discussion

In this study, we engineered the pSS03 binary vector to facilitate high-efficiency genome editing in the Bhimkol banana (*Musa balbisiana*). By leveraging the Cas12a system for the targeted disruption of the *INNER NO OUTER* (*INO*) gene, we established the foundational requirements and expected for seed reduction *via* targeted mutagenesis in this cultivar.

The use of a 23-nt sgRNA of *INO* gene and a T-rich PAM (TTTA) is particularly advantageous for *Musa* genomics for several reasons like AT-Rich targeting in the banana genome, particularly in intergenic and regulatory regions, is characterized by high AT content (Rukavtsova *et al*., 2022). Unlike Cas9, which requires a G-rich (NGGG) PAM, Cas12a’s preference for TTTA motifs allows for much greater flexibility in selecting optimal target sites within the *Musa* B-genome (Tripathi *et al*., 2021). Moreover, Cas12a creates a 5’ overhang (staggered cut) rather than a blunt cut, which is associated with more predictable and often larger deletions that are effective for complete gene knockouts. Research by Malzahn *et al*. (2019) and Ramirez-Torres *et al*. (2021) have shown that Cas12a editing efficiency is significantly enhanced at temperatures between 26-28º C. In the current study, banana tissue culture and embryogenic cell suspension (ECS) protocols typically operate at these temperatures (26-28º C), Cas12a often outperforms Cas9 in tropical fruit crops. The use of the rice (OsU6B) promoter is a well-established standard for driving sgRNA expression in monocots. It ensures that the RNA Polymerase III transcription initiates precisely at the first nucleotide of the guide sequence, which is critical for the secondary structure of the CRISPR complex (Ntui *et al*., 2020).

The methods described for the transformation and regeneration of Bhimkol (*Musa balbisiana*) represent a highly robust protocol that aligns with current “gold standard” practices for banana biotechnology. By integrating Cas12a-mediated editing with an *Agrobacterium*-mediated transformation of Embryogenic Cell Suspensions (ECS), this work overcomes the typical recalcitrance associated with wild banana genomes. The use of *Agrobacterium* strain AGL1 to infect ECS derived from male floral buds of Bhimkol is the most efficient pathway for generating uniform, non-chimeric mutants in banana. AGL1 is a hypervirulent strain known to enhance T-DNA integration in monocots (Mwaka *et al*., 2023). Regenerating plants from a single transformed cell in suspension prevents chimerism, a common issue when using organized tissues like shoot tips (Ntui *et al*., 2020). Previous studies on East African Highland bananas have achieved stable gene integration and high regeneration frequencies using similar ECS-based protocols (Ntui *et al*., 2020). In the current study, we emphasize a high level of stringency by maintaining 40 mg/L hygromycin throughout all stages. This rigorous approach is crucial; in banana transformation, “escapes” (untransformed plants surviving selection) are a frequent hurdle. By keeping selection pressure through maturation (M3), germination (M4), and rooting (R1) stages, it ensured that only truly transgenic events proceed to hardening. The inclusion of 250 mg/L timentin effectively suppresses *Agrobacterium* overgrowth, which can otherwise trigger tissue necrosis or systemic contamination during the sensitive regeneration phase. The timeline described 3 months for embryo maturation followed by 2 months for shoot differentiation is consistent with high-efficiency *Musa* transformation reports (Mwaka *et al*., 2023). The shift to a 16-hour light photoperiod at the M4 stage is a critical physiological trigger for the development of green embryoids, marking the transition from heterotrophic growth to autotrophic potential.

Targeting the second exonic region of the *INO* gene is a deliberate choice intended to ensure a functional knockout of *INO* gene in Bhimkol. In plant biology, the *INO* gene (a YABBY family transcription factor) is essential for the initiation and asymmetric growth of the outer integument of the ovule. Studies in other basal angiosperms, such as *Annona squamosa*, have demonstrated that mutations in the *INO* locus led to seedless phenotypes. By targeting an early exon, the researchers ensure that any resulting INDELs trigger a frameshift mutation, leading to the truncation of essential DNA-binding domains (Stros *et al*.,2007). The molecular characterization of the edited Bhimkol (*Musa balbisiana*) lines confirms the precision of the Cas12a system in generating frameshift mutations led to disruptions of the *INO* gene in Bhimkol. The identification of Type I (1 bp insertion) and Type II (4-7 bp deletions) mutations aligns perfectly with the known repair mechanisms of Cas12a. Unlike Cas9, which usually produces blunt ends, Cas12a creates a 5-nucleotide staggered overhanging end. Research by Zhai *et al*. (2022) noted that this staggered cleavage often leads to larger, more robust deletions through the Non-Homologous End Joining (NHEJ) pathway, as seen in our lines 03_05, 03_06, and 03_17. The single-base (G) insertion in line 03_01 is a documented outcome where the 5’ overhangs are filled in by endogenous DNA polymerases prior to ligation. The frameshift mutations observed in our edited lines are the primary goal when targeting the 2^nd^ exonic regions of *INO* gene to ensure a complete “knockout” of the protein function. Utilizing ICE (Inference of CRISPR Edits) software from Synthego provides a high level of bioinformatic confidence. ICE is widely cited in recent literature (Conant *et al*., 2022) for its ability to accurately quantify editing efficiency and identify specific INDEL distributions from standard Sanger sequencing. This is particularly useful in *Musa* species, where polyploidy or heterozygosity can sometimes cloud manual chromatogram interpretation. A mutation frequency of 23.5% observed in our edited lines is consistent with established CRISPR/Cas12a benchmarks in monocots. While efficiency can vary by target site, previous studies in banana have reported similar ranges (15%–30%) when using Cas12a to target developmental genes (Rukavtsova *et al*., 2022).

The analysis of the Open Reading Frames (ORFs) and predicted amino acid (AA) sequences in the edited Bhimkol lines provides a functional bridge between genetic mutation and biological “knockout.” By demonstrating that the observed INDELs lead to severe protein truncation, this study confirms the successful disruption of the INNER NO OUTER (INO) protein architecture. The wild-type (WT) INO protein in Bhimkol is a 214-AA transcription factor. As a member of the YABBY family, its function is contingent upon two highly conserved structural motifs: the N-terminal zinc-finger domain and the C-terminal YABBY domain (Bowman and Smyth, 1999). The edited Bhimkol lines produced proteins ranging from only 31 to 52 AAs. In line 03_17 (31 AAs) and 03_05 (33 AAs), the truncation occurs so early that neither the zinc-finger nor the YABBY domain is translated. Research by Gross-Hardt *et al*. (2002) on the *INO* locus has shown that the C-terminal YABBY domain is essential for DNA binding and the specification of abaxial cell fate in the outer integument. Without these domains, the protein cannot regulate the downstream genes required for ovule development, effectively mimicking the “seedless” phenotype observed in natural stenospermocarpic varieties (Leon-Reyes *et al*., 2022). The transition of the coding sequence (CDS) from 645 bp to a maximum of 159 bp in the edited lines is the result of Nonsense-Mediated Decay (NMD) precursors. In plant molecular biology, the presence of a premature stop codon (PTC) significantly upstream of the original termination site often triggers the Nonsense-Mediated Decay pathway. According to Shaul (2015), this surveillance mechanism degrades the aberrant mRNA transcripts to prevent the translation of potentially deleterious truncated proteins. The consistency of these PTCs across different edited lines of Bhimkol (e.g., at 159 bp, 102 bp, 105 bp and 96 bp) suggests that the Cas12a-induced mutations are highly effective at ensuring a total loss-of-function (knockout) rather than a mere reduction in activity (knockdown). Similar studies on Cavendish bananas targeting *Phytoene Desaturase* (*PDS*) genes, Naim *et al*., 2018 have shown that early exonic INDELs are the most reliable method for creating stable, heritable knockouts in the *Musa* genus. By using ORF finder tools to confirm the reduction of the protein to less than 25% of its original size, we have provided robust evidence that the pSS03 construct has functionally inactivated the *INO* gene in the Bhimkol (B-genome) background.

The structural annotation of the INO protein *via* the SMART (Simple Modular Architecture Research Tool) platform provides the definitive biochemical “smoking gun” for the functional knockout of the gene in Bhimkol. INO protein consists of two domains, the first domains is HMG_BOX 2 Domain, in wild-type (WT) Bhimkol identified High Mobility Group (HMG) boxes between residues 111 and 165, which are critical for DNA bending and facilitating the assembly of transcriptional complexes. The complete loss of this domain in all edited lines of Bhimkol (03_01, 03_05, 03_06, and 03_17) implies that any truncated peptide produced lacks the machinery required to reorganize chromatin or initiate transcription at target loci (Stros, 2010). The second YABBY domain is a helix-loop-helix motif (spanning residues 9-162 in the WT) essential for DNA binding. Our results showed a reduction of this domain to a mere fraction of its original length (31–51 residues). According to Bowman and Smyth (1999), the integrity of the secondary structures within the YABBY domain is non-negotiable for function; our edited line of Bhimkol effectively delete the core DNA-binding apparatus. In the context of Cas12a-mediated editing, these results highlight the efficiency of targeting the second exon to induce a “polar effect” on protein architecture. Since the INDELs occurred early in the coding sequence (CDS), the downstream HMG_BOX 2 was never translated. This ensures that all essential motifs downstream of the edit site are lost, creating a “structural null” mutant. Research on the *INO* gene in other species, such as *Annona squamosa* and *Arabidopsis*, has shown that even missense mutations within the YABBY domain led to a “seedless” phenotype because the outer integument fails to initiate (Leon-Reyes *et al*., 2022). Our study achieves a more robust knockout by entirely deleting the architectural domains rather than just altering a single amino acid. Utilizing the SMART platform (Letunic and Bork, 2018) added significant rigor to our findings. SMART is widely utilized in plant genomics to predict how INDELs translate into functional changes. By visualizing the shift from a multi-domain regulatory protein (WT) to a short, domain-less peptide (edited lines), we provide clear evidence that the pSS03 construct successfully neutralized the *INO* gene. This domain-level analysis corroborates our ORF data, showing that the genetic frameshift translates directly into a loss of the protein’s “functional toolkit.”

The observation that the edited lines (03_01, 03_05, 03_06, and 03_17) are morphologically similar to the Wild Type (WT) is a significant finding. The *INNER NO OUTER (INO*) gene is primarily involved in the development of the outer integument of the ovule. In *Arabidopsis* and other angiosperms, *INO* expression is highly localized and does not play a role in vegetative growth, leaf morphology, or root development (Leon-Reyes *et al*., 2022). The “normal” vegetative phenotype of the edited lines suggests that our edits have successfully targeted the reproductive trait (seeds) without inducing “off-target” effects or “linkage drag” that often accompanies traditional breeding. Currently, these edited lines are being maintained in a contained transgenic greenhouse. Future phenotyping will be required to confirm the stability of the seedless fruit trait once the plants reach maturity. This is the hallmark of precise genome editing -improving a specific consumer-facing trait (seedlessness) while preserving the agronomic resilience of the wild Bhimkol banana variety.

## Conclusion

In this study, we report the first successful genome editing of the *INO* gene expected Bhimkol, a seedless banana variety (*Musa balbisiana*, BB genome), utilizing the CRISPR/Cas12a system. Our results confirm that CRISPR/Cas12a induces precise modifications, specifically insertions and deletions downstream of the Protospacer Adjacent Motif (PAM) and within the targeted single-guide RNA1 (sgRNA1) regions. This aligns with the nuclease’s established behaviour of producing staggered double-strand breaks. The edited Bhimkol plantlets generated through this research are anticipated to exhibit either reduced seed density or complete seedlessness. These traits will be definitively validated once the plantlets reach reproductive maturity and bear fruit. Our findings establish CRISPR/Cas12a as a highly efficient tool for trait improvement in Bhimkol bananas and other commercially significant crops. This technology provides a streamlined alternative to traditional breeding, which is particularly challenging in polyploid or vegetatively propagated species. The plantlets produced in this study are currently classified as transgenic because they retain the integrated T-DNA from the *Agrobacterium*-mediated delivery system. Consequently, they are subject to existing national biosafety regulations. To facilitate easier commercial adoption and bypass the stringent regulatory hurdles associated with GMOs, the ongoing research will pivot toward: 1) RNP-Based Delivery: Utilizing Ribonucleoprotein (RNP) complexes to achieve “DNA-free” editing. 2) SDN1 Classification: By ensuring the absence of foreign DNA, these plants can be categorized as Site-Directed Nuclease 1 (SDN1). 3) Non-GMO Labelling: Under current Indian biotechnology guidelines, SDN1-edited crops that lack transgene integration can be treated as conventional varieties, significantly accelerating their path to market.

## Acknowledgment

The authors gratefully acknowledge the Department of Biotechnology (DBT), Govt. of India for the primary financial assistance that enabled the successful completion of this study. We acknowledge Department of Agricultural Biotechnology and the DBT-Northeast Centre for Agricultural Biotechnology (DBT-NECAB), Assam Agricultural University, Jorhat, for providing the laboratory facilities and a supportive research environment. We extend our sincere gratitude to Prof. T.W. Okita, whose laboratory provided all the vectors utilized in the Bhimkol research. This research also acknowledges the financial support from ICAR, New Delhi for the ICAR-SRF fellowship provided during the Ph.D. program.

## Author Contributions

SS conceptualized, designed, and supervised the entire study. MNC and AG performed the experiments and analysed the data. MNC and DLS wrote the manuscript with contributions and inputs from all the authors. SKH and TWO critically reviewed and edited the manuscript. All authors have read and approved the final version for publication.

