## Supplementary Fig 1A and 1B for "CRISPR/Cas12a-Mediated Knockout of *INNER NO OUTER (INO)* Gene in *Musa balbisiana* cv. Bhimkol"

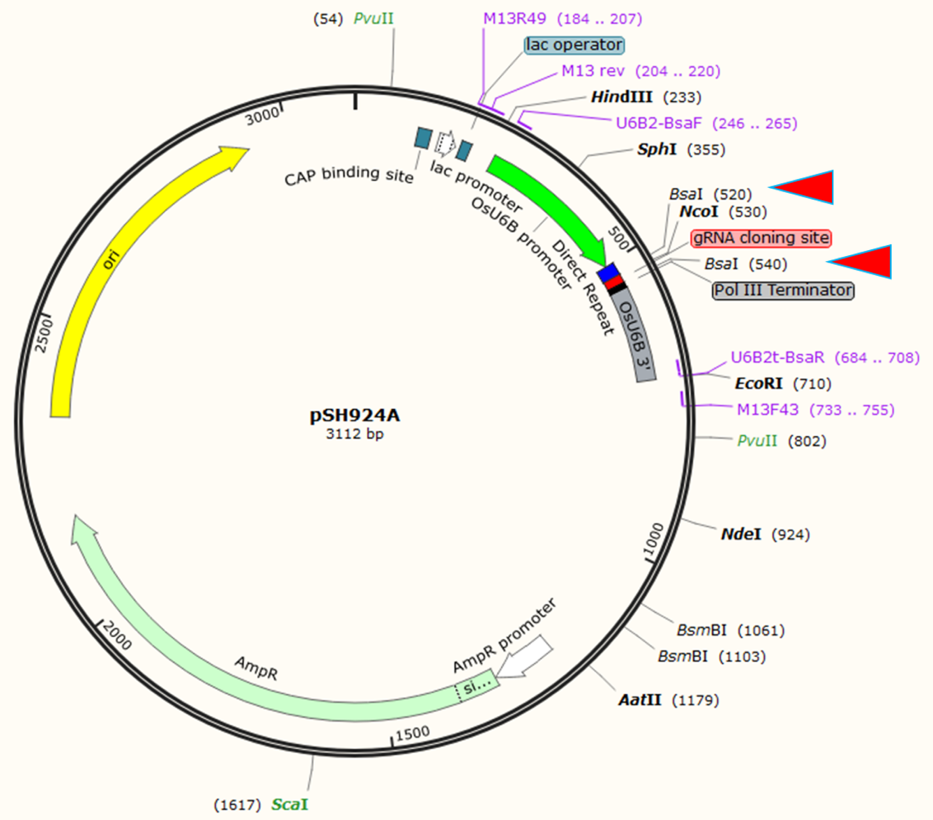

**Supplementary Figure 1A: Physical map of the expression vector- pSH924A**

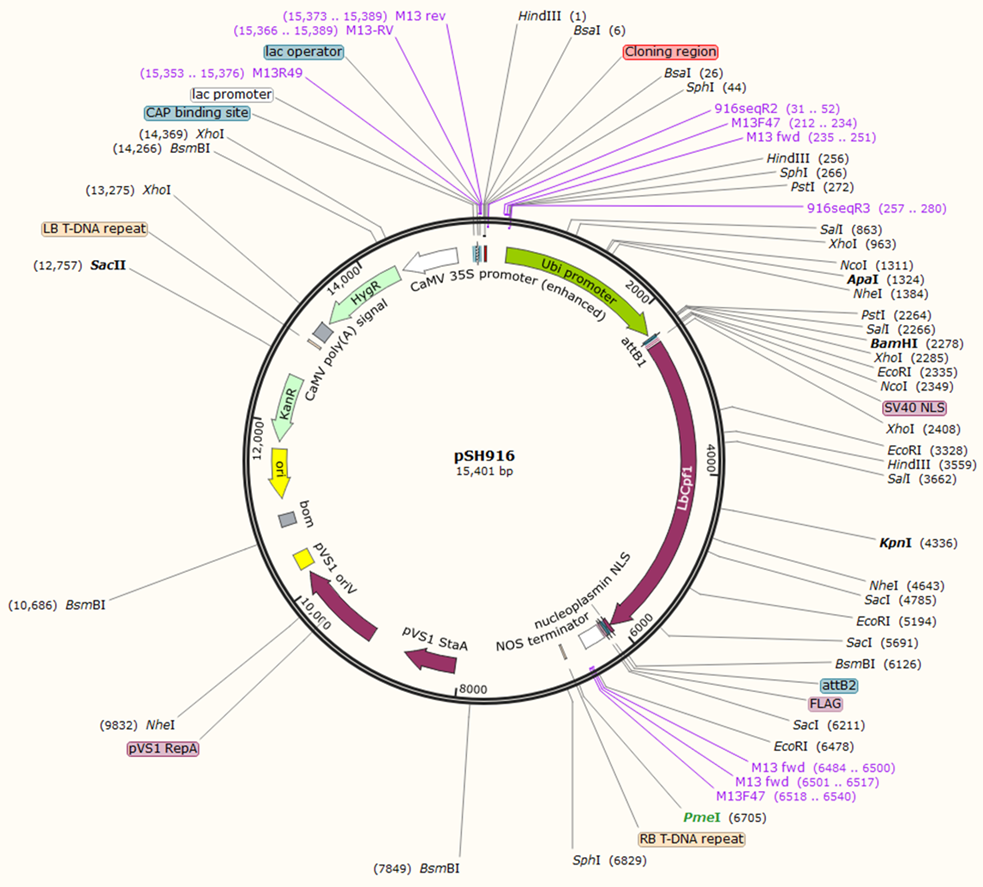

**Supplementary Figure 1B: Physical map of the binary vector – pSH916**
