## Supplementary Fig 2 for "CRISPR/Cas12a-Mediated Knockout of *INNER NO OUTER (INO)* Gene in *Musa balbisiana* cv. Bhimkol"

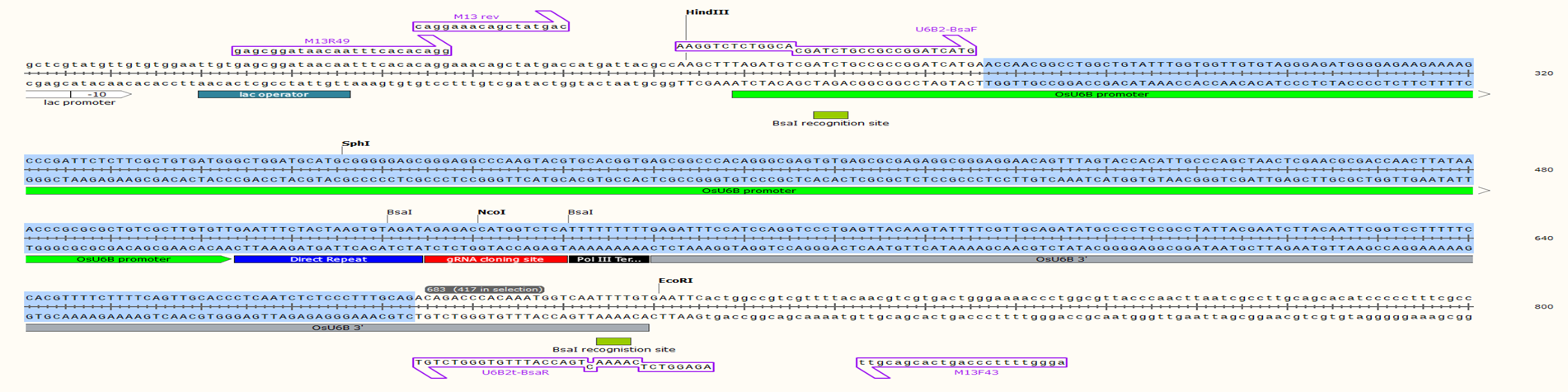
**Supplementary Figure 2:** Position of primers shown for PCR amplification of whole expression cassette U6B2:: INO-sgRNA1::Pol IIIT using U6B2-BsaI F and U6B2t-BsaI R primers.
