## Supplementary Fig 3 for "CRISPR/Cas12a-Mediated Knockout of *INNER NO OUTER (INO)* Gene in *Musa balbisiana* cv. Bhimkol"

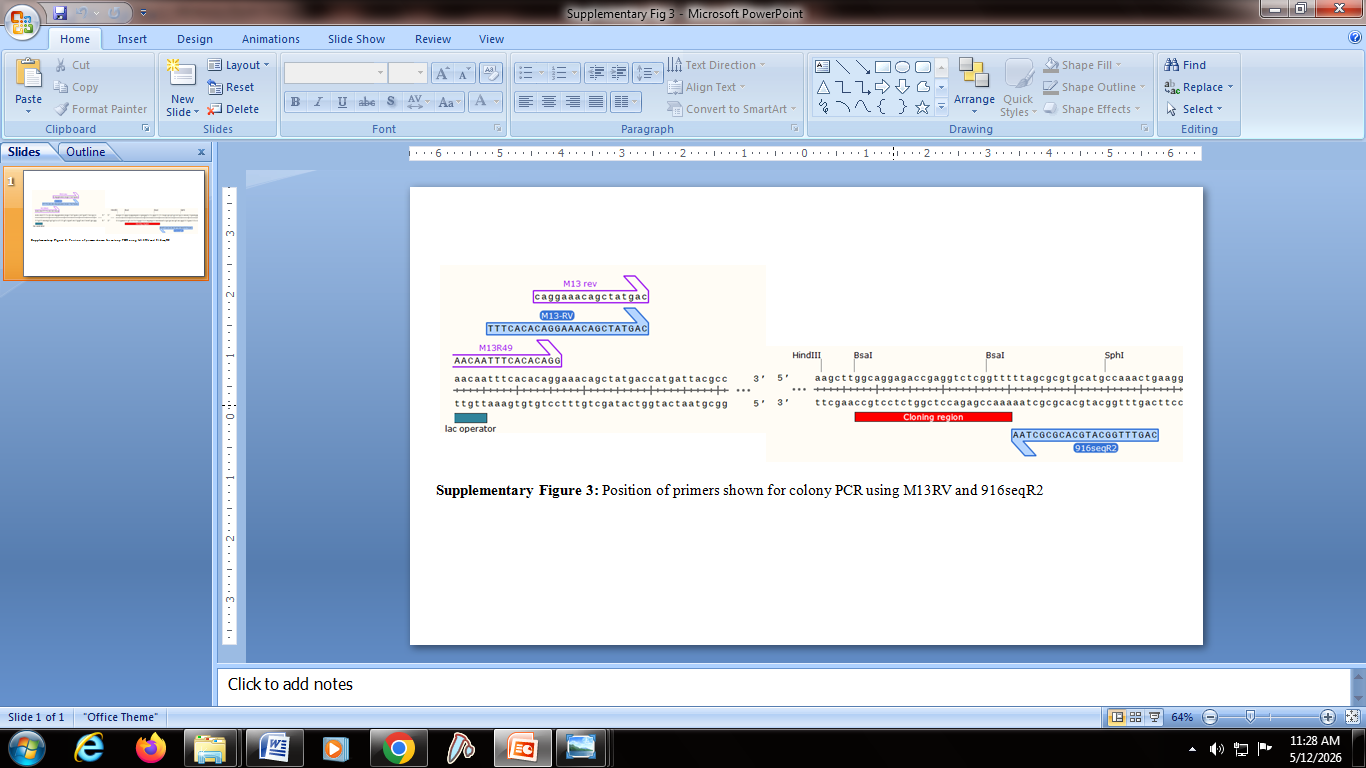

**Supplementary Figure 3:** Position of primers shown for colony PCR using M13RV Forward and 916seqR2 Reverse primers
