## Supplementary tables for "CRISPR/Cas12a-Mediated Knockout of *INNER NO OUTER (INO)* Gene in *Musa balbisiana* cv. Bhimkol"

#### **Supplementary Table 1: Composition of MS (Murashige and Skoog, 1962) stock solutions**

| Component | For 1 litre |
| --- | --- |
| NH4NO3 | 16.5 g |
| KNO3 | 19.0 g |
| CaCl2.2H2O | 4.4 g (Added separately during medium preparation) |
| MgSO4.7H2O | 3.7 g |
| KH2PO4 | 0.85 |

####	**MS- 1 macro (10X concentration)**

**MS-2 Micro (100X** concentration)

| Component | For 1 litre |
| --- | --- |
| MnSO4.H2O | 1690 mg |
| ZnSO4.7H2O | 860 mg |
| H3BO4 | 620 mg |
| KI (1 g/10 ml stock) | 830 µl |
| Na2MO.2H2O (1 g/10 ml stock) | 250 µl |
| CuSO4.5H2O (1 g/10 ml stock) | 25 µl |
| CoCl2.6H2O (1 g/10 ml stock) | 25 µl |

####	**MS-3 vitamins (100X** concentration)

| Component | For 1 litre |
| --- | --- |
| Glycine | 200 mg |
| Nicotinic Acid | 50 mg |
| Pyridoxine HCl (100 mg/l stock) | 500 µl |
| Thiamine HCl (100 mg/l stock) | 100 µl |
| Myo-inositol | 100 mg (Added separately during medium  preparation) |

**MS-4 Fe-EDTA (100X concentration)**

| Component | For 1 litre |
| --- | --- |
| EDTA.2H2O | 3.72 g |
| FeSO4.7H2O | 2.78 g |

**Bluggoe Vitamins (100 X concentration)**

| Component | For 1 litre |
| --- | --- |
| Nicotinic Acid | 50 mg |
| Pyridoxine HCl (100 mg/l stock) | 500 µl |
| Thiamine HCl (100 mg/l stock) | 100 µl |

#### **Supplementary Table 2: Composition of Callus Induction Medium (CIM)**

| Component | For 1 litre |
| --- | --- |
| MS-1 | 100 ml |
| MS-2 | 10 ml |
| MS-3 | 10 ml |
| MS-4 | 10 ml |
| Biotin (1 mg/ml) | 1 ml |
| 2,4-D (4 mg/ml) | 1 ml |
| IAA (1 mg/ml) | 1 ml |
| NAA (1 mg/ml) | 1 ml |
| Sucrose | 30 g |
| Gelling agent (Phytagel) | 2 g |
| pH | 5.7 |

**NB:** Added freshly myo-inositol 100 mg/l and CaCl2 440 mg/l

Added the plant growth hormones like 2, 4- D, NAA and IAA after autoclaving the nutrient medium

#### **Supplementary Table 3: Composition of M2 Suspension medium**

| Component | For 1 litre |
| --- | --- |
| MS-1 | 50 ml |
| MS-2 | 10 ml |
| MS-4 | 5 ml |
| Bluggoe Vitamins | 10 ml |
| Ascorbic acid | 10 mg |
| 2,4-D (1 mg/ml) | 1 ml |
| Zeatin (1 mg/ml) | 250 µl |
| Sucrose | 30g |
| pH | 5.7 |

**NB:**	Added freshly CaCl2 at 220 mg/l.

Added plant growth hormones like 2, 4- D and Zeatin after autoclaving the nutrient medium

#### **Supplementary Table 4: Composition of Co-Cultivation (CCM) medium**

| Component | For 1 litre |
| --- | --- |
| SH salts (Sigma Aldrich) | 3.2 g |
| Sucrose | 45 g |
| Biotin (1 mg/ml) | 1 ml |
| Malt extract | 100 mg |
| Glutamine (100 mg/ml) | 1 ml |
| Ascorbic acid | 10 mg |
| 2,4-D (1 mg/ml) | 1 ml |
| Acetosyringone (200 µM) | 1 ml |
| Phytagel | 2 g |
| pH | 5.3 |

**NB:** SH medium (Shenck and Hilderbrandt, 1972) basal powder from Sigma Aldrich

#### **Supplementary Table 5: Composition of M3 somatic embryo maturation medium**

| Component | For 1 litre |
| --- | --- |
| SH salt (Sigma Aldrich) | 3.2 g |
| Sucrose | 45 g |
| Biotin (1 mg/ml) | 1 ml |
| Malt extract | 100 mg |
| Glutamine (100 mg/ml) | 1 ml |
| Ascorbic acid | 10 mg |
| Picloram (1 mg/ml) | 1 ml |
| Phytagel | 2 g |
| pH | 5.3 |

**NB:** SH medium (Shenck and Hilderbrandt, 1972) basal powder from Sigma Aldrich

Added Picloram after autoclaving the nutrient medium.

Added hygromycin (40 mg/l) for selection of transformed somatic embryos and timentin (250 mg/l) for limiting the overgrowth of *Agrobacterium*

#### **Supplementary Table 6: Composition of Morel and Whitmore Vitamins 100x concentration**

| Component | For 1 litre |
| --- | --- |
| Myo-inositol | 10 g |
| Folic acid | 1 g |
| Nicotinic Acid | 100 mg |
| Pyridoxine HCl (100 mg/ml stock) | 1 ml |
| Thiamine HCl (100 mg/ml stock) | 1 ml |
| Calcium Pentothenate (100 mg/ml stock) | 1 ml |
| Biotin (100 mg/ml stock) | 100 ml |

**Supplementary Table 7: Composition of M4 somatic embryo germination medium**

| Component | For 1 litre |
| --- | --- |
| MS-1 (Macro) | 100 ml |
| MS-2 (Micro) | 10 ml |
| MS-4 (Fe-EDTA) | 10 ml |
| Morel and Whitmore Vitamins | 10 ml |
| Sucrose | 45 g |
| BAP (4 mg/ml) | 1 ml |
| Phytagel | 2 g |
| pH | 5.8 |

**NB:** Added BAP after autoclaving the nutrient medium

Added Hygromycin (40 mg/l) for selection of transformed somatic embryos and

Timentin (250 mg/l) for	limiting the overgrowth of *Agrobacteriu*

#### **Supplementary Table 8: Composition of M5 shoot elongation medium**

| Component | For 1 litre |
| --- | --- |
| MS-1 (Macro) | 100 ml |
| MS-2 (Micro) | 10 ml |
| MS-4 (Fe-EDTA) | 10 ml |
| Morel and Whitmore Vitamins | 10 ml |
| Sucrose | 45 g |
| BAP (2 mg/ml) | 1 ml |
| Phytagel | 2 g |
| pH | 5.8 |

**NB:** Added freshly CaCl2 (440 mg/l)

Added BAP after autoclaving the nutrient medium

Added Hygromycin (40 mg/l) for selection of transformed somatic embryos and Timentin (250 mg/l) for limiting the overgrowth of *Agrobacterium*

#### **Supplementary Table 9: Composition of R1 rooting medium**

| Component | For 1 litre |
| --- | --- |
| MS-1 | 100 ml |
| MS-2 | 10 ml |
| MS-3 | 10 ml |
| MS-4 | 10 ml |
| Sucrose | 30 g |
| Ascorbic acid | 10 mg |
| IBA (2 mg/ml) | 1 ml |
| pH | 5.7 |
| Phytagel | 2g |

**NB:** Added freshly myo-inositol 100 mg/l and CaCl2 440 mg/l

Added the IBA after autoclaving the nutrient medium.

Added Hygromycin (40 mg/l) for selection of transformed somatic embryos and Timentin (250 mg/l) for limiting the overgrowth of *Agrobacterium.*

**Supplementary Table 10: Composition of Mgl Medium**

| Component | For 1 litre |
| --- | --- |
| Mannitol | 100 mg |
| L- Glutamic acid | 1.0 g |
| KH2PO4 | 0.25 g |
| NaCl | 0.1 g |
| MgSO4.7H2O | 0.1 g |
| Tryptone | 5.0 g |
| Yeast extract | 2.5 g |
| pH | 7.0 |
| Bacterial agar | 15.0 g |

#### **Supplementary Table 11: Composition of B5 vitamins**

| Component | For 100 ml |
| --- | --- |
| Myo inositol | 10 g |
| Nicotinic Acid | 100 mg |
| Pyridoxine HCl | 100 mg |
| Thiamine HCl | 1g |

**Supplementary Table 12: Composition of *Agrobacterium* resuspension (AA) medium**

| Component | For 1litre |
| --- | --- |
| N6 basal mixture | 3.98 g |
| B5 Vitamins | 1 ml |
| Sucrose | 68.5 g |
| Glucose | 3.6 g |
| Casein Hydrolysate | 500 mg |
| pH | 5.2 |
| Acetosyringone (100 µM) | 1 ml |

**NB:** N6 medium (Chu, 1975) basal salts from Duchefa Biochemie

Added Acetosyringone after autoclaving the nutreint medium.

### **Supplementary Table 13: Composition of *Agrobacterium* Culture (AB) medium**

| Components | For 1 lit |
| --- | --- |
| Glucose | 5g |
| K_2_HPO_4_ | 3g |
| NaH_2_PO_4_.2H_2_O | 1.3g |
| NH_4_Cl | 150mg |
| CaCl_2_.2H_2_O | 10mg |
| FeSO_4_.7H_2_O | 2.5mg |
| Bacto Agar | 15g |
| pH | 7.2 |
| 1M MgSO_4_.7H_2_O | 1.2ml |

**NB:** Added 1M MgSO_4_.7H_2_O after autoclaving the medium.
